# Transcriptome-based host epistasis and pathogen co-expression in barley-powdery mildew interactions

**DOI:** 10.1101/2023.09.18.558274

**Authors:** Valeria Velásquez-Zapata, Schuyler Smith, Priyanka Surana, Antony V.E. Chapman, Roger P. Wise

## Abstract

*Mildew locus a (Mla*) from the Triticeae grain crop barley (*Hordeum vulgare* L.) encodes a multi-allelic series of nucleotide-binding leucine-rich repeat (NLR) immune receptors. These variable NLRs recognize complementary secreted effectors from the powdery mildew fungus, *Blumeria hordei* (*Bh*), to block disease progression. We used a dynamic time-course transcriptome of barley infected with *Bh* to infer gene effects and epistatic relationships governed by the *Mla6* NLR, two other host loci critical to the interaction, *Blufensin1* (*Bln1*) and *Required for Mla6 resistance3 (Rar3),* and genes that interact with them. *Bln1* is an *R*-gene independent regulator of immunity and the resistant *bln1* mutant exhibits enhanced basal defense to compatible *Bh*. *Rar3* is required for MLA6-mediated generation of H_2_O_2_ and the hypersensitive reaction; the *rar3* mutant contains an in-frame Lys-Leu deletion in the SGT1-specific domain that compromises immunity by a subset of *Mla* alleles. Interactions of *Mla6* and *Bln1* resulted in symmetric, suppression and masked epistasis on the *Bh-*induced barley transcriptome. Likewise, dominant or equal effects were caused by *Mla6* and *Sgt1.* Of a total of 468 barley NLRs, 366 were expressed in our dataset and 115 of those were grouped under different gene effect models, which localized to several chromosome hotspots. The corresponding *Bh* infection transcriptome was classified into nine co-expressed modules, linking differential expression with pathogen development. Expression of most of 517 *Bh* effectors exhibited dependence on disease phenotype and was associated with appressorial or haustorial structures, suggesting that disease is regulated by a host-pathogen intercommunication network that diversifies the response.

## INTRODUCTION

Plant pathogenic fungi are among the greatest deterrents to crop production worldwide (Oerke 2006; Dangl et al. 2013). Obligate biotrophs, such as mildews and rusts, cause some of the most destructive epidemics; however, they are unable to survive autonomously, and as such, present an ideal opportunity to examine interdependent gene regulation between disease agents and their hosts (Dean et al. 2012; Barsoum et al. 2019). Yet, much remains to be discovered regarding the temporal and spatial control of these interconnected processes (Kemen et al. 2011; Salguero-Linares et al. 2022). We have used the association between the powdery mildew fungus, *Blumeria hordei* (*Bh*), and its diploid host plant, barley (*Hordeum vulgare* L.) to tease apart the complex genetics underlying their interaction. Subsequent to germination of haploid conidia and appressorial penetration of the host cuticle, *Bh* secretes effector proteins into epidermal cells via haustorial feeding structures (Kwaaitaal et al. 2017; Jaswal et al. 2020). These effectors allow the fungus to evade host immune responses and acquire nutrients to support colonization (Krattinger and Keller 2016; Lu et al. 2016; Menardo et al. 2017; Bourras et al. 2018; Saur et al. 2019; Langin et al. 2020; Bauer et al. 2021; Crean et al. 2023; Cao et al. 2023). Disease is blocked by immune receptors encoded by host resistance (*R*) genes (Ngou et al. 2022), designated by the prefix *Ml* [for mildew resistance (Wei et al. 2002)].

Intracellular nucleotide-binding leucine-rich repeat (NLR) immune receptors are encoded by some of the largest gene families in plants, with 5408 in the pan-genome of rice (Stein et al. 2018), up to 7780 in the pan-NLRome of wheat (Walkowiak et al. 2020; Sánchez-Martín and Keller 2021), 468 in barley (Li et al. 2021), and 265 in Arabidopsis (Mondragón-Palomino et al. 2017; Wróblewski et al. 2018). These often occur in clusters throughout their respective genomes (Michelmore and Meyers 1998; van Wersch and Li 2019), and once activated and translated, interact directly or indirectly with effector proteins secreted by pathogens (Borrelli et al. 2018; Ngou et al. 2022). The barley MLA (NLR-type) immune receptor family and its orthologs specify resistance to many cereal diseases, including powdery mildew, stem- and stripe rust, and rice blast (Jordan et al. 2011; Maekawa et al. 2012; Periyannan et al. 2013; Mago et al. 2015; Cesari et al. 2016; Bettgenhaeuser et al. 2021; Brabham et al. 2022). In some cases, the identical MLA host protein determines recognition specificity to effectors from evolutionary diverged pathogens, for example, the adapted ascomycete *Bh* vs. the non-adapted basidiomycete, *Puccinia graminis striformis* (Bettgenhaeuser et al. 2021), or *Bh* and *Magnaporthe grisea*, the causal agent of rice blast (Brabham et al. 2022).

To facilitate their function, host NLR immune receptors are often stabilized by additional proteins, for example, heat shock protein 90 (HSP90), required for *Mla12* resistance1 (RAR1), and suppressor of the G2 allele of SKP1 (SGT1) (Shen et al. 2003; Bieri et al. 2004; Halterman and Wise 2004; Shirasu 2009; Chapman et al. 2021; Chapman et al. 2022) as well as interacting with and activating each other (Cesari et al. 2014; Wróblewski et al. 2018; Bentham et al. 2020; Sun et al. 2020; Van-Wersch et al. 2020; Ngou et al. 2022). On the pathogen side, coding sequences (CDS) for 534 candidate secreted effector proteins (CSEPs) have been identified in the *Bh* genome (Pedersen et al. 2012; Menardo et al. 2017; Frantzeskakis et al. 2018). These were annotated by being predominantly expressed in the haustorium, with a predicted signal peptide, no transmembrane domain and no or little homology outside the powdery mildews (Spanu et al. 2010; Pedersen et al. 2012; Kusch et al. 2014; Frantzeskakis et al. 2018). A subset of these have been functionally characterized by identification of their host targets and/or gene silencing (Zhang et al. 2012; Pliego et al. 2013; Ahmed et al. 2015; Aguilar et al. 2016; Pennington et al. 2019; Yuan et al. 2021; Li et al. 2022; Velásquez-Zapata et al. 2023).

Gene effect models can explain the genetic interactions that underlie host disease responses. Of these, one of the most utilized involves the estimation of epistasis, where gene effects are classified as additive or the product of gene interaction(s) (Mani et al. 2008; Dixon et al. 2009). When single mutants of two genes are associated with a phenotype whose sum differs from the double mutant, this non-additivity indicates that epistasis is present (the two genes interact genetically). In this regard, RNA sequencing (RNA-Seq) can be used as a proxy for gene expression as a quantitative phenotype to perform gene effect analyses (Angeles-Albores et al. 2018). Gene expression is also heritable and linked to phenotypic variation at the microscopic and macroscopic levels (Blanc et al. 2021). Here, we used a dynamic transcriptome collected from the interaction between barley and *Bh,* to explore gene effects, epistatic relationships, and co-expression networks governed by the NLR-type *R*-gene, *Mla6*, and two additional host genes critical to the interaction, *Required for Mla6 resistance 3* (*Rar3*) (Chapman et al. 2021), and *Blufensin1* (*Bln1*) (Meng et al. 2009; Xu et al. 2015; Guo et al. 2022). Two epistasis models were inferred in the host; and in parallel, temporally associated co-expression modules were distinguished in the pathogen. Genomic location is proposed as a possible mechanism of the host gene effects by associating chromosome hotspots with the different genetic effect patterns. Results from this analysis point to a large perturbation network of host and pathogen arsenals under different genetic mechanisms that diversify expression patterns and increase robustness of the response.

## MATERIALS AND METHODS

### Transcriptome data collection

The temporal transcriptome used for this study was obtained from an infection time course of five barley genotypes with *Blumeria hordei* (*Bh*) isolate 5874 which carries the *AVR_a6_* effector. The barley genotypes consisted of a resistant, wild-type plant (CI 16151) that carries the *Mla6* gene and four fast-neutron-derived mutants. Plants that carry the *bln1* mutation (*bln1*-m19089) are resistant, while the other three are susceptible; an *mla6* mutant (*mla6*-m18982), a *rar3* mutant *(rar3-*m11526), and the double mutant (*mla6*+*bln1*)-m19028. The split-plot design contained 90 samples, comprised of 5 genotypes, 6 timepoints [0, 16, 20, 24, 32, and 48 Hours after Inoculation (HAI)], and 3 three biological replicates. First leaves were challenged with *Bh* isolate 5874 (Caldo et al. 2004) and then leaves were collected at each timepoint, for each genotype and replicate. RNA was extracted with hot (60°C) phenol/guanidine thiocyanate (Caldo et al. 2004; Caldo et al. 2006). Libraries were prepared using the Illumina TruSeq stranded RNA sample preparation kit (Illumina, Inc., San Diego, CA) and sequenced with the Illumina HiSeq2500 system, using a randomized block design with single-end 100-bp reads (**Figure S1**).

### Transcriptome profiling

We utilized the Nextflow DSL2 (Di Tommaso et al. 2017) nf-core/rnaseq pipeline version 3.4 (Ewels et al. 2020; Patel et al. 2021) to align and quantify mRNA sequencing data from barley, *Hordeum vulgare* L., and *Bh*. The 90 RNA-Seq libraries were deposited to NCBI-GEO system, accession GSE101304. The reads were aligned and quantified against the *Bh* DH14 genome assembly (NCBI accession GCA_900239735.1) and annotation from the Ensembl database (https://fungi.ensembl.org/Blumeria_graminis/Info/Index) obtained from Stefan Kusch at RWTH Aachen University (Frantzeskakis et al. 2018). The pipeline for *Bh* was run with the additional parameters: “--skip_multiqc”. Similarly, all RNA-Seq libraries were aligned against the barley Morex v3 reference genome and annotation (Mascher et al. 2021) obtained from the USDA GrainGenes database (Blake et al. 2019). The barley pipeline was run with the following parameters:

> “--skip_multiqc --bam_csi_index --rseqc_modules
>
> ‘bam_stat,infer_experiment,junction_annotation,junction_saturation,read_duplication’”.

The pipeline consisted of adapter and quality trimming using Trim Galore (Andrews 2010; Martin 2011; Krueger 2012), read mapping with STAR (Dobin et al. 2013) and quantification with Salmon (Patro et al. 2017).

### Differential expression analysis

Read counts were normalized using the taxon-specific method (Klingenberg and Meinicke 2017). Each raw count table (barley and *Bh)* was taken and size factors calculated using median-of-ratios normalization (Anders and Huber 2010). Then, normalized tables were combined and used as input for DESeq2 (Love et al. 2014), assigning the calculated size factors to the counts before calculating differentially expressed genes (DEGs). Continuing with the DESeq2 pipeline, DEGs were identified using a model with read counts as response variable, and timepoint*genotype terms as explanatory variables. Results from this analysis, including fold changes and p-values were taken for subsequent analyses. Adjusted p-values (Benjamini and Hochberg 1995) were used to determine DEGs using a value of <0.001 for barley and <0.003 for *Bh*.

### Mutant characterization

Fast-neutron mutants of *Mla6* and *Bln1* were identified as described in Chapman et al. 2021, and distinguished by genetic complementation followed by differential expression, first by Affymetrix Barley1 GeneChip analysis (Meng et al. 2009; Xu et al. 2014), and more recently, by RNA-Seq read mapping between wild-type progenitor and each single mutant (**Figure S2**). The *rar3* mutation was previously characterized (Chapman et al. 2021), consisting of an in-frame Lys-Leu deletion in the SGS domain of Suppressor of the G2 allele of SKP1 (*Sgt1_ΔKL308-309_)*.

To map the extent of the *mla6* and *bln1* mutations, several regions from the barley v3 genome (Mascher et al. 2021) were extracted as follows: 1) We analyzed *Mla* (HORVU.MOREX.r3.1HG0012670) and flanking genes in its proximity that had differential expression between the wild-type and the *mla6-*m18982 mutant; we also included all the genes in the *Mla* locus as reported by (Wei et al. 2002); lastly, all the genes without mapped reads in *mla6-*m18982 were added to the list. 2) *Bln1* was not annotated in the barley v3 genome assembly. To identify the *bln1* mutation, we used BLAST with the *Bln1* sequence (NCBI ID FJ156737.1) against the v3 barley genome to identify its coordinates and then extracted differential expressed genes between wild-type and *bln1*-m19089 mutant in the *Bln1* vicinity. In addition, we added all the genes without reads mapped in the *bln1*-m19089 mutant. To all these lists we added 1000 bp at the 5’ and 3’ ends.

The genomic coordinates were used to extract the corresponding sequences using bedtools (Quinlan and Hall 2010). After extracting the regions of the candidate genes, we mapped the RNA-Seq reads back to them from the wild-type, single and double mutants for the timepoints 16 and 20 HAI. We used BWA-mem (Li and Durbin 2009) for mapping and IGV (Robinson et al. 2011) for visualization. The bam files were used for a reference-guided assembly using Trinity (Grabherr et al. 2011). We compared the mappings and assemblies to identify differences between wild-type progenitor and mutants. Consensus gene candidates across samples were analyzed to identify the effects of the mutations.

### Gene effect models

We designed two methods for evaluating the gene effects of the *Mla6, Bln1* and *Sgt1* genes. A full description of both models is presented in **S1 text**.

### Model *Mla6, Bln1*

The first model (*Mla6, Bln1*) is a full epistasis design as we had data from single and double mutants. Let E(G) be the log of the global gene expression levels of a plant with the genotype G. For analyzing the effects of the pair *Mla6, Bln1* we write the global expression equation for each genotype. If *DB* is defined as the Defense Background genotype, the collection of genes in G involved in defense responses and common to all the genotypes:

> E*(Mla6, Bln1) = Mla6 + Bln1 + Mla6*Bln1 + Mla6*DB + Bln1*DB + DB*

> E*(Mla6, bln1) = Mla6 + Mla6*bln1 + Mla6*DB + DB = Mla6 + Mla6*DB + DB*

> E*(mla6, Bln1) = Bln1 + Bln1*DB + DB*

> E*(mla6, bln1) = DB*

We can calculate differentially expressed genes among a pair of genotypes (DEG(G1/G2)) as the difference between their expression E(G1) *–* E(G2). For example, DEGs between CI 16151 (WT) and each of the mutants:

> DEG*(Mla6, Bln1/Mla6, bln1) = E(Mla6, Bln1) – E(Mla6, bln1) = Bln1 + Mla6*Bln1 + Bln1*DB*

> DEG*(Mla6, Bln1/mla6, Bln1) = E(Mla6, Bln1) – E(mla6, Bln1) = Mla6 + Mla6*Bln1 + Mla6*DB*

> DEG*(Mla6, Bln1/mla6, bln1) = Mla6 + Bln1 + Mla6*Bln1 + Mla6*DB + Bln1*DB*

And with the double mutant as reference:

> DEG*(Mla6, bln1/mla6, bln1) = Mla6 + Mla6*DB*

> DEG*(mla6, Bln1/mla6, bln1) = Bln1 + Bln1*DB*

> DEG*(Mla6, Bln1/mla6, bln1) = Mla6 + Bln1 + Mla6*Bln1 + Mla6*DB + Bln1*DB*

We assigned DEG()=0 as a fold change equal to zero, *i.e.*, there are no significant differences. DEG()*≠*0 means there are significant differences and considering all values of the fold change. DEG()>0, DEG()<0 indicate significant differences and the sign of the fold change in the comparison. A change in the fold change proportion larger than 20% was considered significant. Genes whose expression deviated from the expected value were classified as epistatic while the ones that fitted the expected value were considered additive.

1. Epistatic effects, Epi(*Mla6, Bln1*):

> DEG(*Mla6, Bln1/mla6, bln1*) – DEG(*Mla6, Bln1/mla6, Bln1*) – DEG(*Mla6, Bln1/Mla6, bln1*)*≠0*
2. No epistasis, additive effects of *Mla6* and *Bln1*, Addi(*Mla6, Bln1):*

> DEG(*Mla6,* Bln1*/mla6, bln1*) – DEG(*Mla6, Bln1/mla6, Bln1*) – DEG(*Mla6, Bln1/Mla6, bln1*)*=0*
3. Symmetric epistasis:

> Epi(*Mla6, Bln1*) ∩ [DEG(*Mla6, bln1*/*mla6, bln1*) = 0 ∩ DEG(*mla6, Bln1*/*mla6, bln1*) = 0]
4. Masked epistasis:

> Epi(*Mla6, Bln1*) ∩ [DEG(*Mla6, bln1*/*mla6, bln1*) *≠* 0 ∩ DEG(*mla6, Bln1*/*mla6, bln1*) = 0]
5. Suppression epistasis:

> Epi(*Mla6, Bln1*) ∩ [DEG(*Mla6, bln1*/*mla6, bln1*) *=* 0 ∩ DEG(*mla6, Bln1*/*mla6, bln1*) *≠* 0]
6. Pseudo Masked epistasis:

> Epi(Mla6*, Bln1*) ∩ [DEG(*Mla6, bln1* /*mla6, bln1*) > 0 ∩ DEG(*mla6, Bln1* /*mla6, bln1*) < 0] *OR*

> Epi(*Mla6, Bln1*) ∩ [DEG(*Mla6, bln1* /*mla6, bln1*) < 0 ∩ DEG(*mla6, Bln1* /*mla6, bln1*) > 0]
7. Positive epistasis:

> Epi(*Mla6, Bln1*) ∩ [DEG(*Mla6, bln1* /*mla6, bln1*) < 0 ∩ DEG(*mla6, Bln1*/*mla6, bln1*) < 0]
8. Negative epistasis:

> Epi(*Mla6, Bln1*) ∩ [DEG(*Mla6, bln1*/*mla6, bln1*) > 0 ∩ DEG(*mla6, Bln1* /*mla6, bln1*) > 0]

### Model *Mla6, Sgt1*

The second model (*Mla6, Sgt1*) was defined taking the wild-type expression as reference, since the double mutant was not available. This model consisted in separating the gene effects by comparing the DEG lists from the comparison of the *mla6* and *rar3* (*Sgt1_ΔKL308-309_)* single mutants with CI 16151 (WT). The intersection DEG(*Mla6, Sgt1*/*Mla6, sgt1*) ∩ DEG(*Mla6, Sgt1*/*mla6, Sgt1*) contains the shared effects between *Mla6* and *Sgt1.* Then, the difference between the DEG of each wild-type : mutant comparison contains the dominant effects of each gene. Now let’s compute each of the described lists:

1. Shared effects between *Mla6* and *Sgt1:* Shared(*Mla6, Sgt1)=*

> DEG(*Mla6, Sgt1*/*Mla6, Sgt1_ΔKL308-309_*)*≠0* ∩ DEG(*Mla6, Sgt1*/*mla6, Sgt1*)*≠0*
2. Dominant effects of *Sgt1:*

> Dom(*Sgt1*)= [DEG(*Mla6, Sgt1*/*Mla6, sgt1*) *≠* 0] – Shared(*Mla6, Sgt1)*
3. Dominant effects of *Mla6:*

> Dom(*Mla6*)= [DEG(*Mla6, Sgt1*/*mla6, Sgt1*) *≠* 0] – Shared(*Mla6, Sgt1)* This intersection DEG(*Mla6, Sgt1*/*Mla6, Sgt1_ΔKL308-309_*) ∩ DEG(*Mla6, Sgt1*/*mla6, Sgt1*) can be split in the genes that have an equal effect between the two mutants (with the same fold-change and no significant differences when they are compared), or predominant effects when the wild-type version of each gene (*Mla6* or *Sgt1*) has a greater effect on gene expression:
4. Equal effects between *Mla6* and *Sgt1:*

> Equal(*Mla6, Sgt1) =* Shared(*Mla6, Sgt1)* ∩ [DEG(*mla6, Sgt1*/*Mla6, Sgt1_ΔKL308-309_*) = 0]
5. Predominant effects of *Sgt1:*

> PreDom(*Sgt1*) *=* Shared(*Mla6, Sgt1)* ∩ [DEG(*mla6, Sgt1*/*Mla6, Sgt1_ΔKL308-309_*)>0]
6. Predominant effects of *Mla6:*

> PreDom(*Mla6*) *=* Shared(*Mla6, Sgt1)* ∩ [DEG(*mla6, Sgt1*/*Mla6, Sgt1_ΔKL308-309_*)<0]

### GO analysis

Genes classified under each gene effect model and per timepoint and their consensus were analyzed using GO enrichment. Using clusterProfiler (Yu et al. 2012) and the GO terms reported for the barley and *Bh* annotations (Frantzeskakis et al. 2018; Mascher et al. 2021) we calculated gene enrichment with a p-value threshold of 0.05. We used ggplot2 (Wickham 2016) to graph the functions of the genes under each pattern over time.

### Chromosomal position analysis

Gene lists under the consensus of each gene effect classification model were used to perform genomic position analysis. Using GenomicRanges (Lawrence et al. 2013), we binned the genes in the barley genome and then we designed a hypergeometric test to look for regions in the genome with enrichment in each gene list. Taking genome windows of 1 Mb, 10 Mb, 100 Mb and the complete chromosome or scaffold we tested for enrichment of genes classified in each of the categories for each model. We then adjusted the p-values using the Benjamini and Hochberg method (Benjamini and Hochberg 1995) and considered significant enrichment using a threshold of 0.05. Barley genes that were classified in each pattern were plotted for the significant regions using the ggbio package (Yin et al. 2012).

### Nucleotide-binding leucine-rich repeat (NLR) pattern analysis

The Morex v3 genome-wide barley NLR list (Li et al. 2021) was taken to study the gene effects of *Mla6*, *Bln1* and *Sgt1*. Gene expression was clustered and a heatmap constructed for each group adding two layers of color annotations (one for each gene effect model). Gene expression patterns for each gene were also generated, calculating significant differences across time for each genotype comparison. These results were summarized using the package multcompView (Graves et al. 2019), and using a letter code to show differences. Patterns were manually curated, and examples shown to characterize the dataset for each of the gene effect models.

### Bh infection kinetics

Microscopic quantification of fungal structures was performed as described in (Chapman et al. 2021). Structures included attached conidiospores, appressorial attachment, and hyphal indices 1, 2 and 3, which are used to estimate % elongating secondary hyphae (ESH) as an indicator of functional haustoria (Ellingboe 1972). Ten-centimeter (cm) seedlings for each timepoint and genotype were assayed in triplicate. Subsequent to inoculation with *Bh* 5874, five leaves assigned to each timepoint were harvested and submerged in clearing solution (3:1, alcohol: acetic acid) for at least 24 h and then transferred to 70% ethanol for another 24 hours and then to 20% ethanol. Scoring was performed by treating the leaves with Coomassie blue for two minutes to visualize spores and hyphae. After staining, leaves were trimmed to 5 cm from the tip and structures were counted on both the abaxial and adaxial sides (**S1 Data**).

### Co-expression analysis

*Bh* normalized gene counts using DESeq2 (Love et al. 2014) were taken for all the replicates, genotypes and timepoints. Genes with complete gene counts were used to build the co-expression network by following the tutorials presented in the Weighted Gene Co-expression Network Analysis (WGCNA) R package (Langfelder and Horvath 2008). The clustering was checked for quality and then co-expression clusters were calculated. To characterize these clusters functionally, we performed GO analysis using a hypergeometric test. GO annotations were obtained using Interproscan (Jones et al. 2014) and functional annotation for the *Bh* genome provided through personal communication with Carsten Pedersen and Hans Thordal-Christensen at the University of Copenhagen. GO analysis was performed for the *Bh* co-expression clusters and enrichment terms (adjusted p-value <0.05) were reported.

The co-expression network was correlated with different phenotype traits including timepoint, disease phenotype and the infection kinetic structures. Significant associations were calculated at the cluster and gene level. Significant associations were characterized by GO analysis. In the case of effectors, a further functional table was obtained to perform the hypergeometric tests (Pedersen et al. 2012).

## RESULTS

### A barley-Bh transcriptome to explore the genetic interactions of Mla6, Bln1 and Sgt1

A 6-point infection time course was conducted via inoculation of barley with *Bh* isolate 5874 (*AVR_a6_*), spanning 0 to 48 hours after inoculation (HAI) and covering *Bh* appressorium formation (0-16 HAI), penetration of epidermal cells (16-20 HAI), and development of haustorial feeding structures (32-48 HAI) (Chapman et al. 2021). As shown in **Figure 1A**, five genotypes were included in the design. The wild-type progenitor, CI 16151, contains the *Mla6* NLR gene, which confers resistance to *Bh* isolates that secrete the AVR_A6_ effector. The susceptible *mla6*-m18982 mutant contains 168 single-nucleotide polymorphisms (SNPs) and 2 x 3-nucleotide indels in the *Mla6* coding sequence, resulting in 85 amino acid changes (**Figure S2 B,C**). *Bln1 is* an *R*-gene independent regulator of immunity, whose silencing results in the down-regulation of transcripts involved in nuclear import and the secretory pathway (Meng et al. 2009; Xu et al. 2015; Guo et al. 2022). The resistant *bln1* mutant, *bln1*-m19089, contains a deletion of the gene (**Figure S2 E**) and exhibits enhanced defense to compatible *Bh* isolates. The susceptible double mutant, *(mla6+bln1)*-m19028, contains both the *bln1* and *mla6* variants. Lastly, *Rar3* is required for *Mla6*-mediated hypersensitive response and generation of H_2_O_2_. The susceptible *rar3-*m11526 *(Mla6, Bln1, Sgt1_ΔKL308-309_)* mutant contains an in-frame Lys-Leu deletion in the SGS domain of the co-chaperone SGT1, which interacts physically with MLA (Shen et al. 2003; Chapman et al. 2021; Chapman et al. 2022).

**Figure 1.**
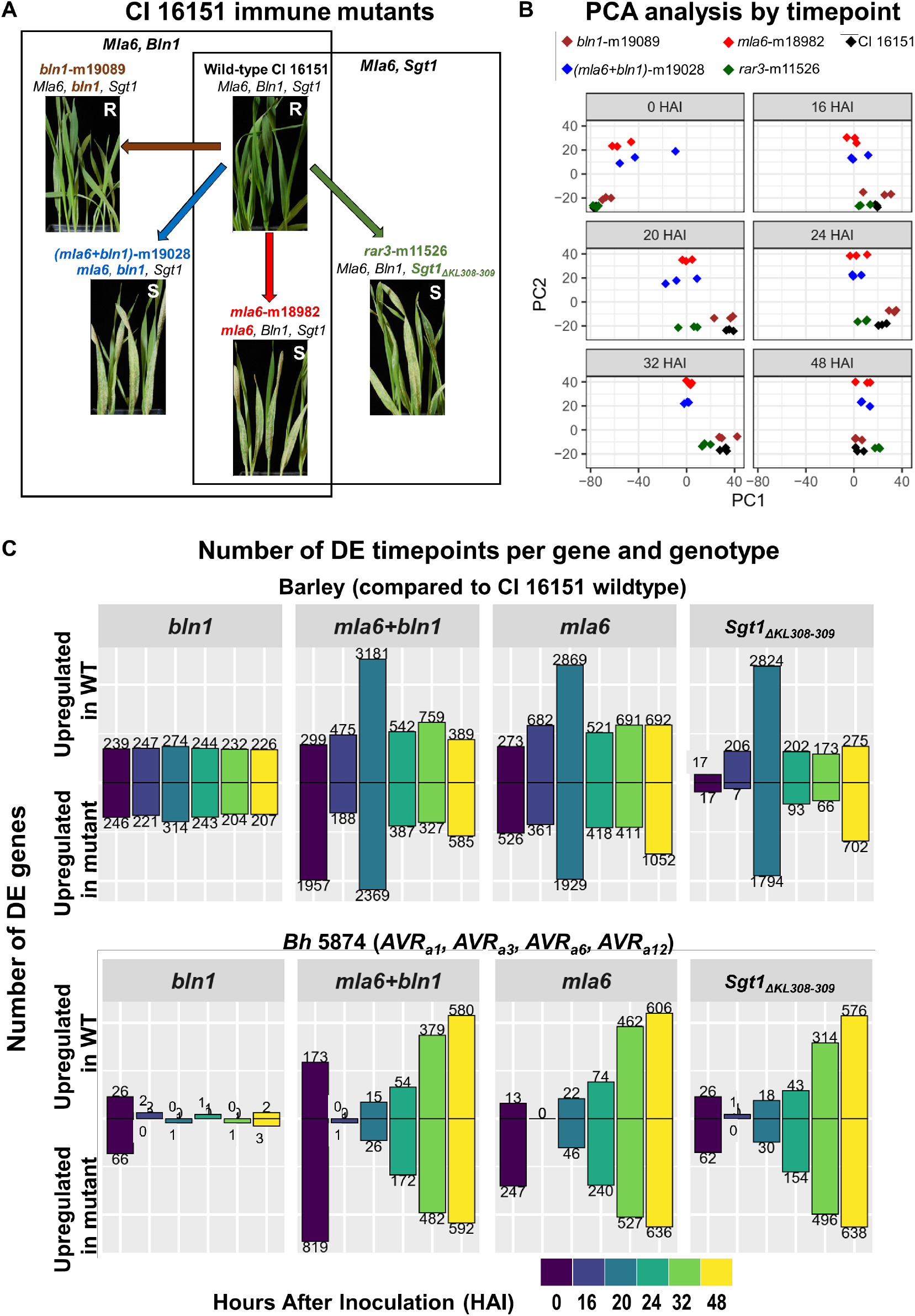
A dynamic barley-*Bh* transcriptome of immune mutants. A) Wild-type progenitor CI 16151, carrying the *Mla6* gene conferring resistance to *AVR_a6_ Bh* isolates, and four fast-neutron derived immune mutants were used to investigate the effects of the functionally connected, but physically unlinked, *Mla6*, *Bln1* and *Sgt1*. Lineages used in each of the *Mla6, Bln1* and *Mla6, Sgt1* models are boxed. B) PCA analysis of the RNA-Seq samples obtained from the immune mutant panel, colored by genotype, and separated by timepoint. C) Number of DE timepoints per gene when the wild-type and each mutant are compared, separated by genotype and organism. DE genes calculated at significance adjusted p-values of 0.001 for barley and 0.003 for *Bh*.

First leaf seedlings inoculated with fresh *Bh* conidiospores were sampled at 0, 16, 20, 24, 32, and 48 HAI (5 genotypes x 6 timepoints x 3 biological replications). Bulk RNA-Seq reads from both barley and *Bh* were processed, and transcript counts analyzed to characterize gene expression. As shown in **Figure 1B**, principle component analysis (PCA) indicated that samples clustered by timepoint and genotype. The initial time point (0 HAI) is separated from the rest in the PC1 axis, which comprises 43% of the variance. The PC2 component contained 16% of the variance. Two genotype groups were identified: *mla6*-m18982 and double mutant *(mla6+bln1)-* m19028 for the first; and CI 16151 (*Mla6, Bln1, Sgt1)*, *bln1-*m19089 and *rar3-*m11526 for the second. Interestingly, the genotype groups did not correspond to disease phenotype, having both resistant and susceptible phenotypes in the second group. When the PCA was separated by barley and *Bh* (**Figure S3**) we could observe that the *Bh* dataset is divided by disease phenotype, a pattern that is reinforced at later timepoints.

Differentially expressed genes (DEGs) were identified per timepoint taking as reference the wild-type progenitor line CI 16151 and comparing across mutant genotypes (**S2 Data**). **Figure 1C** contains a bar plot of the number of DE genes per timepoint in barley or *Bh,* separated by mutant and foldchange (overexpressed in wild-type or in the mutant). We found that all the susceptible mutants showed a peak in the number of barley DEGs at *Bh* penetration (20 HAI), while the *(mla6+bln1)-*m19028 double mutant had the highest number of DE genes/timepoint and the *bln1*-m19089 single mutant had the lowest. Lastly, when focusing on the resistant *bln1*-m19089 mutant, we observed a low number of DEGs across the time course, without a peak as the susceptible mutants (a range 433-588 DEGs per timepoint).

*Bh* displayed different expression patterns from those found in the host. Most *Bh* DEGs were associated with the development of haustoria (32 and 48 HAI), starting with almost no DEGs at penetration of barley epidermal cells (16 HAI) and gradually increasing over time. We also observed that the number of *Bh* DEGs in the resistant *bln1*-m19089 mutant were almost non-existent across the time course. These observations indicate that the host environment perceived by *Bh* and specifically the plant disease phenotype is the main determinant of *Bh* gene expression.

### Global and gene-wise epistasis indicate that Mla6 and Bln1 interact genetically

Our mutant panel enabled us to define a model of epistasis between *Mla6* and *Bln1* to study their effects on barley transcriptome dynamics. Using gene expression as phenotype, we calculated the genome-wide epistatic effects of *Mla6* and *Bln1,* using a global index as described by Angeles-Albores and colleagues (2018). Predicted additive effects of these genes were calculated as the fold change by comparing each single mutant, *mla6*-m18982 or *bln1-* m19089, to their progenitor, CI 16151, using DESeq2 (Love et al. 2014). Observed genetic effects would correspond then to the fold change of the comparison between the double mutant, (*mla6*+*bln1*)*-*m19028, and the CI 16151 genotypes (see Methods). **Figure 2A** shows the genome-wide comparison between the predicted additive and the deviation from the observed genetic effects of *Mla6* and *Bln1*, colored by timepoint. From this plot we can estimate the global epistasis index as the slope of the line. According to this model, a slope of zero indicates no epistasis, and deviations from zero indicate global epistasis. Subsequently, we calculated the range at each timepoint, obtaining values of -0.78 to -0.62. These values (different than zero) indicate symmetric epistasis between *Mla6* and *Bln1*, and evidence of stronger genetic effects of *Mla6* (negative slope values) on barley gene expression.

**Figure 2.**
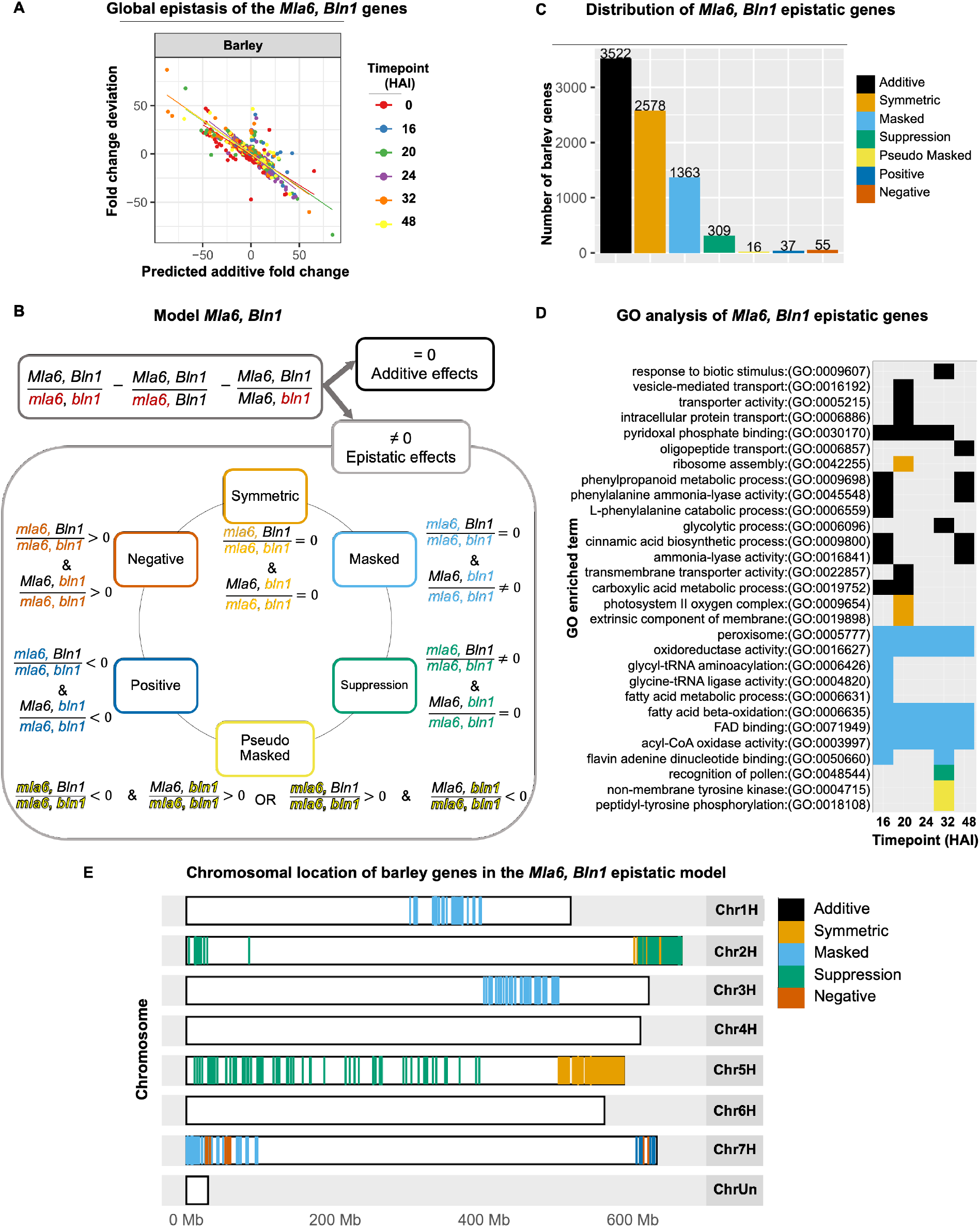
Global and gene-wise epistasis between *Mla6* and *Bln1.* A) Global epistasis between *Mla6* and *Bln1* in the *Bh-*induced barley transcriptome. The expected additive fold change between the single and double mutant and the observed deviation is plotted and separated by organism and timepoint. B) Proposed gene-wise epistasis model between *Mla6* and *Bln1* applied to the *Bh-*induced barley transcriptome. Additive and epistasis effects are separated and then the epistatic effects are classified in six categories. C) Distribution of the *Mla6* and *Bln1* epistatic classifications for the *Bh-*induced barley transcriptome. D) Gene Ontology (GO) enrichment analysis of the barley epistatic patterns applied to the transcriptome across time. E) Enriched chromosomal locations of the barley genes under the *Mla6, Bln1* epistatic model. Each barley chromosome is shown, and colored areas correspond to the different genetic patterns significantly enriched (adjusted p-value<0.05).

To explore the transcriptional effects of *Mla6* and *Bln1* at the gene-level we applied an epistasis classification model (**S1 Text**). As illustrated in **Figure 2B**, the model includes additive and interaction effects and those patterns with non-additive effects are further classified into six categories (Cordell 2002; Dixon et al. 2009). These include symmetric epistasis, where both single and double mutants have the same effect; masked and suppression epistasis where the expression of the double mutant is equal to one of the single mutants (*mla6*-m18982 and *bln1-* m19089, respectively); pseudo masked epistasis, where the expression of the double mutant is between the two single mutants. Two additional classes were found where the double mutant has a higher (positive epistasis) or lower (negative epistasis) expression than the single mutants.

These classifications were applied to our RNA-Seq dataset, separating by timepoint. To generate a consensus of the epistasis patterns across time, we used the most frequent classification across all timepoints for each gene (excepting 0 HAI), and then summarized the results as shown in **Figure 2C** (full list in **S3 Data**). We determined that the 0 HAI timepoint introduced error into the consensus because of artificially low read counts, resulting in large differences observed in the PCA analysis with other timepoints (**Figure 1B**) (Gihawi et al. 2023). Subsequently, the ensuing dataset would have a higher false positive rate for mining DEGs. Most of the barley genes had an additive effect between *Mla6* and *Bln1*, which means they did not go through epistasis. Among the epistatic genes we found that most had a symmetric pattern followed by masking and suppression. These results coincide with what we observed in the genome-wide epistasis analysis presented in **Figure 2A**. Smaller sets of genes presented pseudo masked, positive and negative epistasis.

We then performed Gene Ontology (GO) analysis to describe the biological functions of the genes under each epistasis pattern, as shown in **Figure 2D**. Additive effects of *Mla6* and *Bln1* targeted genes involved in intracellular and membrane transport, MAP and receptor kinase, response to stimulus, chloroplast, and carbohydrate binding. These additive effects were found across the entire time course, with higher frequency at 16 and 48 HAI, which are associated with *Bh* penetration and formation of haustoria, respectively. Among epistatic patterns, symmetric epistasis was associated with genes at 20 HAI, which are involved in ribosomal, photosynthetic, and extrinsic component of membrane functions. Pseudo-masked and suppression epistasis influenced genes at 32 HAI, which were involved in phosphorylation and recognition of pollen. Lastly, masked epistasis presented an influence across the time course, and the genes under this pattern were associated with oxidoreductase activity, peroxisome, translation, fatty acid and acyl CoA metabolism.

After finding evidence of genetic interaction between *Mla6* and *Bln1*, we explored possible mechanisms of regulation of such patterns. We looked for association between the genes under each pattern and their genomic location (see Methods). Taking genome windows of 1 Mb, 10 Mb, 100 Mb and the complete chromosome or scaffold we tested for enrichment of genes classified in each of the categories for each model. We found significant genomic hotspots associated with each epistasis category as shown in **Figure 2E** and **S4 Data**, distributed across chromosomes 1H-3H and 5H-7H. Genes under symmetric epistasis were concentrated in the chromosomal regions 2H.6, 3H.5, 5H.5 and 6H.0, of 100 Mb in size. Masked epistasis had effects on genes concentrated in the regions 1H.0 and 7H.0 while genes under suppression were associated with the regions 2H.0, 2H.6 and to a wide range in chromosome 5 from 5H.0 to 5H.3. Lastly, genes under negative epistasis were enriched to regions in chromosome 7 (7H.0 and 7H.6). The distribution of the hotspots was diverse, showing high location specificity with two exceptions: region 2H.6 and 7H.0 that were enriched for two epistatic patterns.

### Design of a Mla6, Sgt1 gene effect model

We proposed a second classification model for gene expression associated with *Mla6* and *Sgt1* which did not require a double-mutant dataset. However, we do know that *Mla6* requires *Sgt1* to function properly, *i.e.*, express resistance to *AVR_a6_* containing *Bh* isolates (Chapman et al. 2021; 2022). For this second model (*Mla6, Sgt1*), we could not separate additive from epistatic effects, but we could compare the strength of the contribution of each gene to the wild-type genotype. To accomplish this, we used a classification of five categories by separating the effects of *Mla6* and *Sgt1* into dominant, predominant, and equal effects (**Figure 3A**). We defined dominant effects as those where only one of the single mutants is differentially expressed as compared to the wild-type. For example, genes under dominant effects of *Mla6* were calculated as those that were not DE between the wild-type and *rar3-*m11526, and DE when the wild-type was compared to *mla6-*m18982. Then, we further separated the shared effects (genes that are DE in both single mutants), into predominant and equal effects. Predominant effects have significantly higher expression in one of the single mutants as compared to the other one. We defined genes under predominant effects of *Mla6* as those whose expression was higher in *rar3-*m11526 as compared to *mla6-*m18982, as the first contains a wild-type version of the gene, indicating its stronger effect. Lastly, equal effects of *Mla6* and *Sgt1* have the same expression level in both mutants.

**Figure 3.**
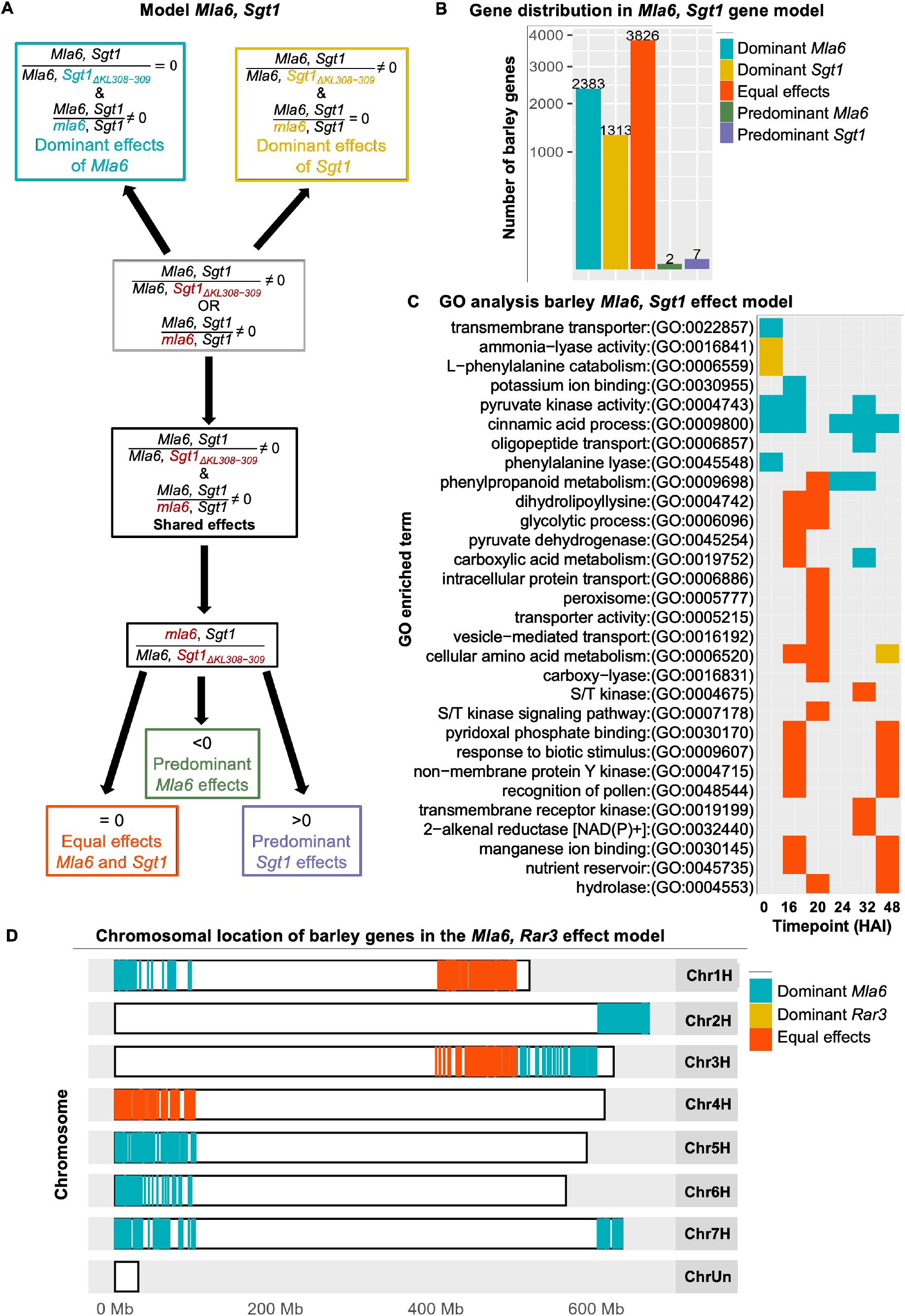
Genetic effects model of *Mla6* and *Sgt1*. A) Proposed effect model between *Mla6* and *Sgt1* in the *Bh-*induced barley transcriptome. Effects are separated in dominant effects where only one of the mutant genotypes has significant differences; equal effects were both mutants have the same effects and predominant effects where the strength of the differences are higher in one of the mutants. B) Distribution and C) GO enrichment across time of the model classifications for the *Bh-*induced barley transcriptome. D) Enriched chromosomal locations of the barley genes under the *Mla6, Sgt1* epistatic model. Each barley chromosome is shown, and colored areas correspond to the significantly enriched genetic patterns (adjusted p-value<0.05).

Figure 3B shows the distribution of these classifications for the barley transcriptome and the full list is reported in **S3 Data**. Results indicate that most of the DE genes in the barley transcriptome were under equal gene effects of *Mla6* and *Sgt1*, followed by those under the dominant effects of *Mla6* and of *Sgt1.* GO enrichment analysis of the barley genes (Figure 3C) associated with each classification showed that the expression of genes under the same effects of *Mla6* and *Sgt1* were involved in diverse biological processes across the time course. Among them we found serine/threonine and tyrosine kinase, intracellular and vesicle−mediated transport, peroxisome, pyruvate metabolism, phenylalanine ammonia−lyase, response to biotic stimulus and cinnamic acid biosynthesis. Dominant effects of *Mla6* were associated with the expression of genes involved in transmembrane transporter, potassium ion binding, cinnamic acid process, phenylalanine lyase and phenylpropanoid metabolism. Dominant effects of *Sgt1* were associated with the expression of genes with functions such as L−phenylalanine catabolism and cellular amino acid metabolism.

When the *Mla6*, *Sgt1* classification model was analyzed for associations with genomic location, we found enrichment of genes associated with dominant effects of *Mla6* and equal effects of *Mla6* and *Sgt1* (**S4 Data**). Genes associated with dominant effects of *Mla6* were enriched to several regions across six out of seven barley chromosomes (Figure 3D) including 1H.0, 2H.6, 3H.5, 5H.0, 6H.0, 7H.0 and 7H.6. Genes whose expression was equally affected by *Mla6* and *Sgt1* were enriched to three chromosomal regions: 1H.4, 3H.4 and 4H.0. These results indicate that the hotspots associated with dominant effects of *Mla6* were shared with the epistatic patterns in the *Mla6*, *Bln1* model while novel locations were enriched when genetic effects of *Mla6* and *Sgt1* were analyzed. For example, regions 1H.4, 3H.5 and chromosome 4 were only enriched in genes under this type of classification. These results indicate different pathways of each interaction with *Mla6*.

### Epistasis regulates NLR expression

We explored the effects of epistasis on transcript accumulation that encode NLRs, key proteins that determine the outcome of barley-*Bh* interactions. The 468 NLRs annotated for the TRITEX v3 assembly (Li et al. 2021) were filtered under each gene effect model. Of these, 366 were present in our expression dataset and 115 were classified under at least one of the models (103 for the *Mla6, Bln1* epistasis model and 90 for the *Mla6, Sgt1* gene effect model). Figure 4 depicts a heatmap for these 115 NLRs across time and genotypes. Using hierarchical clustering we grouped the NLRs and annotated them using the two gene effect models. First, for the *Mla6, Bln1* model we observed that the most common pattern within the NLR group was additive effects followed by symmetric and masked epistasis. Forty-four NLRs were under additive effects and were scattered across the heatmap with four subgroups of more than two members which presented co-expression. The additive epistasis group included NLR genes with very diverse expression patterns. *Mla6* (r3.1HG0012670) belongs to this group, presenting a flat expression pattern in the *mla6* single and *mla6 + bln1* double mutant.

**Figure 4.**
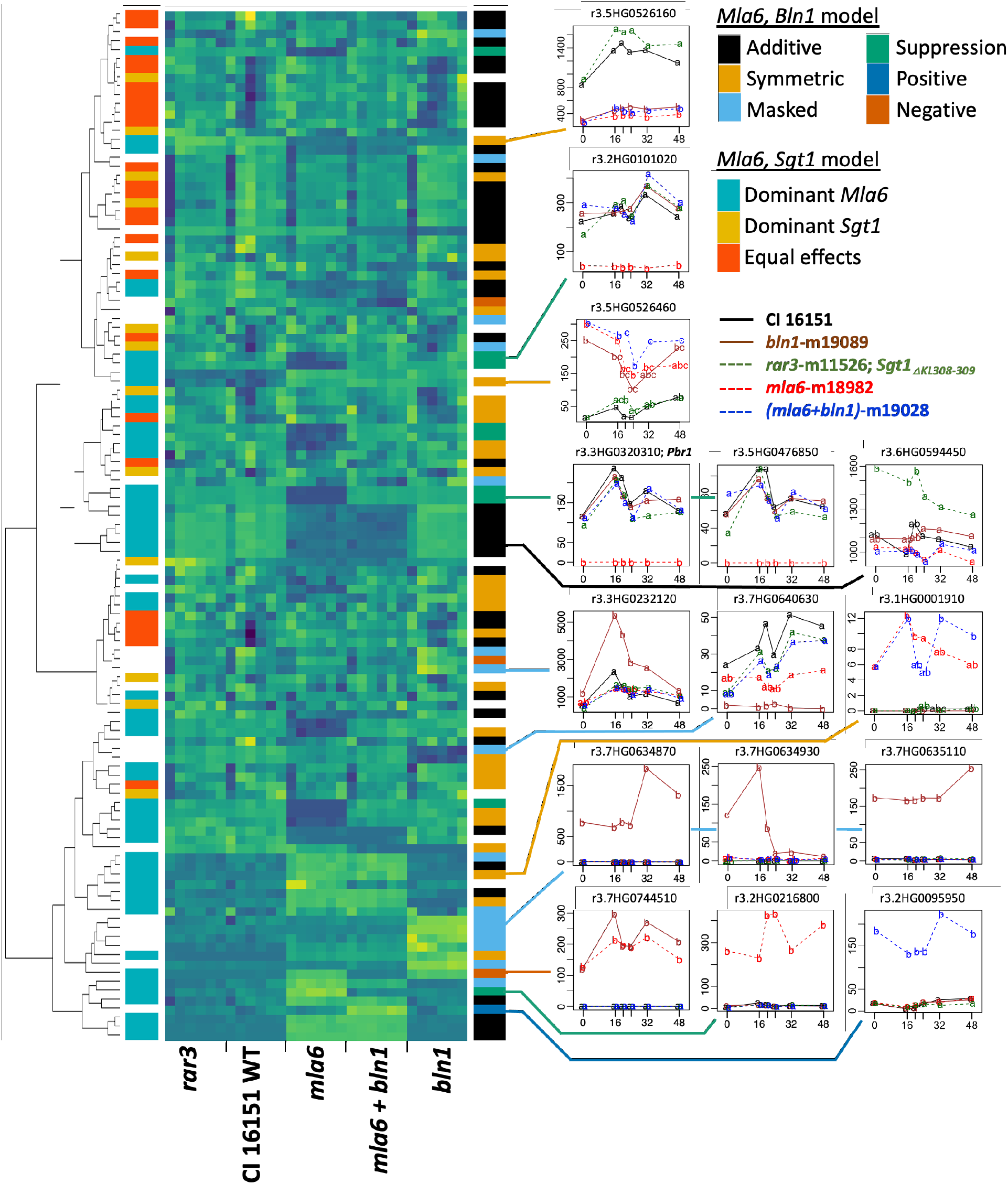
Heatmap of the barley NLRs classified in the *Mla6, Bln1* or *Mla6, Sgt1* gene effect models. Barley NLRs under each model were taken and their expression across genotypes and time were hierarchically clustered and plotted with a heatmap. Annotation of each model is shown at the right and left of the heatmap and. Lastly, examples from each group were plotted with expression patterns as indicated by legend color. An adjusted p-value cutoff of 0.001 was used to assign significant differences in expression.

Similarly, symmetric epistasis was observed in 30 NLR, with a diverse set of expression patterns. We highlight the examples as r3.5HG0526160, where plants harboring single and double mutants of *mla6* and *bln1* were down-regulated as compared to *rar3* or CI 16151; or r3.5HG0526460 with the opposite pattern. Masked epistasis (16 NLRs) had one co-expressed subgroup including representative genes such as r3.7HG0640630 where *mla6* and *bln1* single mutants had intermediate expression, or where in r3.7HG0634870 only the *bln1* mutant had expression counts above zero. Lastly, we also found some NLRs whose expression pattern fit suppression (9 NLRs), positive (1 NLR) and negative (3 NLRs) epistasis. Most genes under suppression epistasis showed no significant transcript accumulation in the *mla6* single mutant, including NLRs such as *Pbr1* (Carter et al. 2019; Jaiswal et al. 2023); in contrast, r3.2HG0216800 was not expressed in the all other genotypes, except the *mla6* single mutant. One NLR presented positive epistasis (r3.2HG0095950) which did not show significant transcript accumulation except in the (*mla6*+*bln1*)*-*m19028 double mutant. Three NLRs presented negative epistasis (r3.7HG0744510, r3.6HG0629180, r3.7HG0744530) and their expression patterns showed higher expression in the single mutants as compared to the double mutant in the dataset.

When the *Mla6, Sgt1* model classifications were analyzed we found that the clustering separated the dominant effects of *Mla6* and equal effects while dominant effects of *Sgt1* were more scattered. We also noticed that dominant effects of *Sgt1* coincided with symmetric epistasis in the *Mla6, Bln1* model. Among them we highlight *Rp1*-like r3.6HG0594450, an NLR whose expression is significantly higher for *rar3-*m11526 *(Mla6, Bln1, Sgt1_ΔKL308-309_)* over the rest of the genotypes. Equal effects of *Mla6* and *Sgt1* were associated with expression patterns where the susceptible genotypes had low transcript levels while the resistant genotypes presented peaks across the time course. In contrast, the group of NLRs that were classified under dominant effects of *Mla6* presented expression patterns where *rar3*-m11526 had a similar expression to the resistant genotypes while the *mla6*-m18982 mutant was significantly different from them, grouping with the double mutant or separating from all the other genotypes.

### Co-expression networks reveal associations between Bh effectors and fungal developmental stages

In contrast to the host, changes in *Bh* expression appear to be due to differences in the environment that the fungus perceives, rather than an epistatic effect from the barley genes. Considering that the *Bh* genotype is constant in the experiment, changes in its transcriptome would be the result of the interaction, which could be either compatible (susceptible for host, virulent for pathogen) or incompatible (resistant for host, avirulent for pathogen). As we observe in Figure 1, the *Bh-*infection transcriptome is associated with the disease phenotype, which led us to select a co-expression network as the modeling method to explore the pathogen response. This analysis enables the association of the gene expression patterns in clusters and their functional association with phenotypic traits (in this case, infection structures over time). The *Bh* co-expression network was constructed using WGCNA (Langfelder and Horvath 2008) using gene counts for all 90 samples and replicates. We identified nine clusters in the network, designated *Bh*1 (black; 135 genes), *Bh*2 (blue; 768 genes), *Bh*3 (brown; 725 genes), *Bh*4 (green; 420 genes), *Bh*5 (grey; 2091 genes), *Bh*6 (pink; 125 genes), *Bh*7 (red; 245 genes), *Bh*8 (turquoise; 1921 genes), and *Bh*9 (yellow; 613 genes). For functional characterization, we performed GO analysis using Interproscan (Jones et al. 2014) and completed an enrichment test for each cluster, as shown in **Figure S4**.

Among the enriched GO terms per cluster, we found RNA binding for *Bh*3 (brown); oxidoreductase and lipid metabolism for *Bh*4 (green); protein kinase and nuclease activity for *Bh*5 (grey); transcription for *Bh*6 (pink), proteolysis and ribosomal for *Bh*8 (turquoise), and catalytic activity for *Bh*9 (yellow). Genes in the co-expression network were then correlated to quantitative differences in fungal development by WGCNA (Langfelder and Horvath 2008). As documented in **S1 Data**, traits comprised microscopic quantification of fungal structures including attached conidiospores, appressorium attachment, and hyphal indexes 1, 2 and 3; observed at 16, 20, 24, 28, 32 and 48 HAI (Chapman et al. 2021).

At the infection kinetics level, significant differences were only found when resistant vs. susceptible genotypes were compared (Chapman et al. 2021), indicating that the *Bh* phenotype follows a similar response as the transcriptome. However, as shown in **S5 Data**, significant associations between the co-expression clusters and the phenotypic traits were identified at the gene level. Significant associations were found between the infection kinetics traits and the modules. First, spore and appressorium attachment had multiple associations with several modules, with opposite correlation signs), while the associations with hyphal indexes were more gene specific. *Bh*5 (gray) and *Bh*9 (yellow) clusters were exclusively associated with spore and appressorium attachments, while *Bh*1 (black), *Bh*3 (brown), *Bh*7 (red), and *Bh*8 (turquoise), were associated with most of the stages.

GO analysis of the genes and clusters associated with each *Bh* stage is shown in Figure 5A. The number of spores across the time course was associated with actin binding and catalytic activity for *Bh*9 (yellow), translation, protein folding and ribosome for *Bh*8 (turquoise), and RNA binding and translation for *Bh*3 (brown). Appressorium attachments had associations with actin binding for *Bh*9 (yellow), transcription factor activity for *Bh*6 (pink), translation and RNA processing for *Bh*3 (brown) and metallopeptidase activity for *Bh*4 (green). Hyphal index 1 was associated with ATP hydrolysis coupled proton transport for *Bh*8 (turquoise). Hyphal indexes 2 and 3 had common associations with *Bh*8 (turquoise), including endopeptidase and oxidoreductase activity, proteasome, and proton transmembrane transporter activity. In addition, hyphal indexes had associations with transported activity for *Bh7* (red) and redox activity for *Bh*4 (green). Hyphal index 2 was also associated with GTP binding and GTPase activity for the *Bh*4 (green), and ribonuclease activity for *Bh*3 (brown).

**Figure 5.**
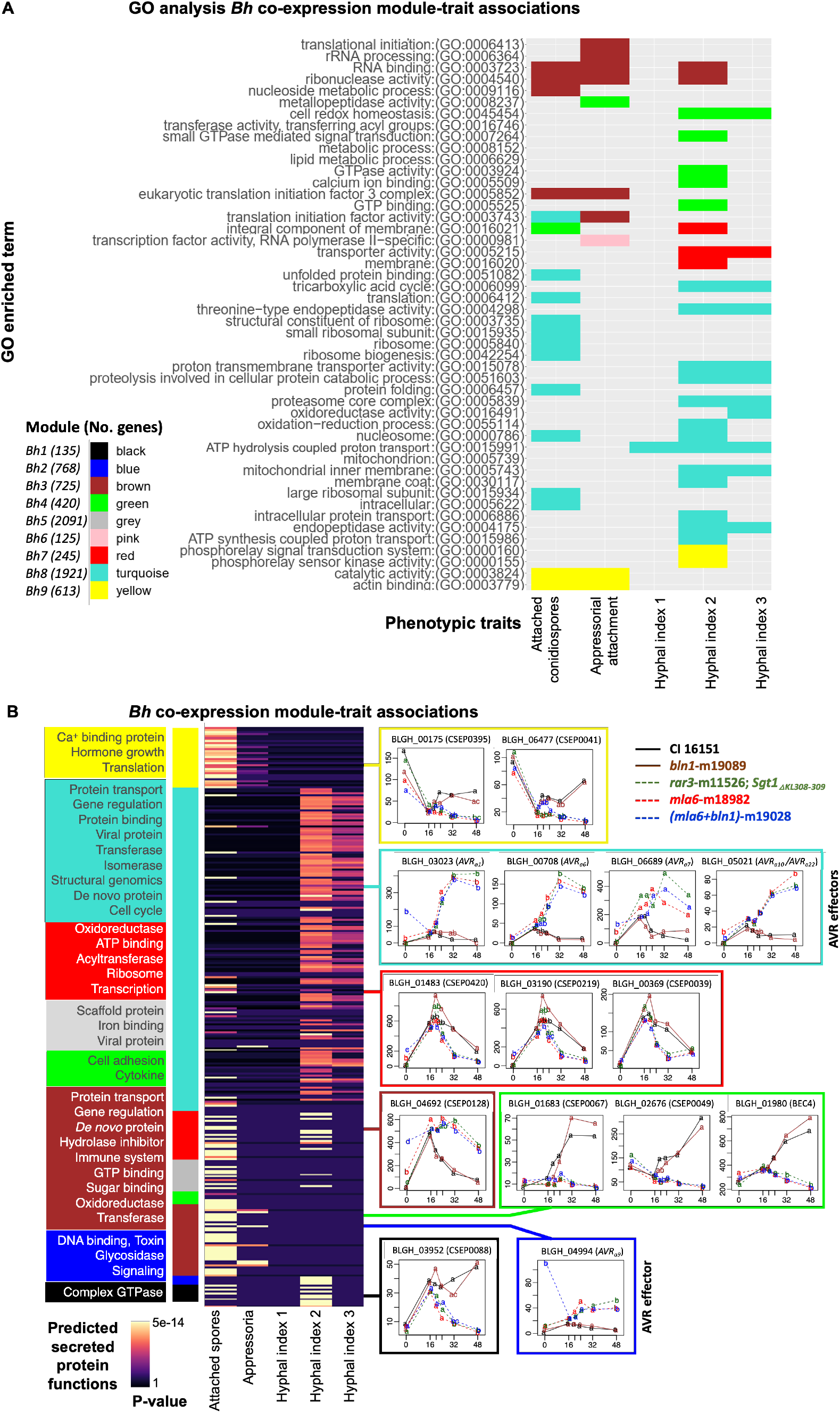
The *Bh* co-expression network is associated with fungal developmental stages. A) GO enrichment analysis of the significant module-trait associations of the *Bh* transcriptome co-expression network. Modules associated with phenotypic traits including developmental stages (spores, appressorium attachment, and hyphal indexes 1-3). B) Significantly associated effectors with phenotypic traits, color represents p-value in the hypergeometric test. On the left, functional prediction of the effectors in each module; on the right, expression patterns of the representative effectors per module. An adjusted p-value cutoff of 0.003 was used to assign significant differences in expression.

Effector proteins secreted by pathogens modulate, inhibit, or accelerate host processes to enable nutrient acquisition and colonization (Lo-Presti et al. 2015; Toruño et al. 2016). From the *Bh* list containing 893 effectors/secreted proteins, as reported by (Pedersen et al. 2012; Frantzeskakis et al. 2018), 864 were expressed in our dataset. We characterized these secreted proteins using predicted functions from (Pedersen et al. 2012; Frantzeskakis et al. 2018) and performing hypergeometric tests for enrichment in each of the modules. The modules that were enriched with secreted proteins included *Bh*7 (red), *Bh*8 (turquoise) and *Bh*3 (brown). The *Bh*7 (red) module was enriched in predicted effector functions including proteasome, periplasmic binding protein, ribosome, and transcription. In the *Bh*8 (turquoise) module we found enrichment for protein transport, gene regulation and protein binding, while effectors in the *Bh*3 (brown) module were associated with gene regulation, *de novo* protein, hydrolase inhibitor and immune system.

When effector expression patterns were analyzed, we found they were associated with disease phenotype, with a separation between resistant vs. susceptible genotypes. We explored the associations between secreted proteins with the phenotypic traits and characterized them functionally for their role during infection. Figure 5B and **S5 Data** contains the enriched functions and associations with phenotypic traits. Here, we highlight effector expression patterns starting with AVRs, *e.g*., BLGH_00708 [identified as *AVR_a6_* and representing a group of effectors from the *Bh*8 module (turquoise)], which is significantly associated with attached spores, and hyphal index 1, 2, and 3. Its expression is significantly higher in susceptible backgrounds after penetration as compared to resistant hosts. Other AVR effectors that fit this expression pattern include BLGH_03023 (*AVR_a1_*; associated with spores, hyphal index 1,2,3), BLGH_05021 (*AVR_a10_/AVR_a22_*; associated with spores, hyphal index 1,2,3), and BLGH_06689 (*AVR_a7_*; associated with spores). BLGH_04994 (*AVR_a9_*) was classified in the *Bh2* (blue) cluster with a similar expression pattern, but was not associated with any infection kinetics traits.

Among the effectors classified from the *Bh*9 (yellow) module we identified BLGH_00175 (CSEP0395) and BLGH_06477 (CSEP0041) as associated with spores and appressorial attachments; these have lower expression after penetration in susceptible as compared to resistant genotypes. Lastly, we found an expression pattern consistent with a peak at penetration followed by a drop in either the resistant or the susceptible background, with significant differences for timepoints after the peak. Examples of this pattern include BLGH_01483 (CSEP0420), BLGH_03190 (CSEP0219), and BLGH_00369 (CSEP0039) from the *Bh*7 (red) module and associated with spores and/or hyphal index 2 and 3.

## DISCUSSION

### Epistasis and NLR expression

Barley is a model large-genome cereal to investigate the genetic mechanisms of disease resistance and susceptibility (Dangl et al. 2013; Mascher et al. 2021). Powdery mildew, as an obligate fungal biotroph, also is an archetype for the intricate interactions with the host and its interconnected transcriptional networks (Spanu et al. 2010; Pedersen et al. 2012; Frantzeskakis et al. 2018). Quantitative transcriptome data can be used to evaluate a wide variety of biological questions, beyond conventional differential expression analysis. We utilized our barley immune mutant collection, all derived from a single progenitor, to develop a custom analysis to interrogate the interactions between barley and powdery mildew from the perspective of gene effect models and co-expression.

The first (*Mla6, Bln1)* gene effect model leveraged single and double mutants of the NLR-type resistance gene *Mla6* (Halterman et al. 2001; Wei et al. 2002) and the cysteine-rich, Ca2+ influx reducing peptide encoded by *Bln1* (Meng et al. 2009; Xu et al. 2015; Guo et al. 2022). In this case, we evaluated epistasis with both a global and a gene-wise estimation. The two analyses were consistent, though, the gene-wise model facilitated identification of specific epistasis patterns per gene and the subsequent functional classification and chromosomal location of each set.

The *Bh*-induced barley transcriptome is mostly under additive effects (no epistasis) of *Mla6* and *Bln1.* The next most abundant pattern corresponds to symmetric epistasis, which classifies those genes whose expression is influenced equally by *Mla6* and *Bln1*, and are involved in ETI and PTI responses, respectively. Suppression- and masked-epistasis were particularly intriguing in terms of their effects on immunity. In suppression epistasis, gene expression is affected by the *mla6* mutation only in combination with the wild-type *Bln1* gene. Generally, most barley genes under suppression epistasis are downregulated in the *mla6* mutant, when compared with the *mla6 + bln1* double mutant (Figure 4) and are primarily clustered at the telomeric ends of chromosome 2H, and the top half of chromosome 5H (Figure 2E). This is exemplified by the barley NLR, AvrPphB Response 1 (HvPBR1), which specifies recognition to the effector AvrPphB from *Pseudomonas syringae* pv. *phaseolicola* (Carter et al. 2019). Co-expression of full-length HvPBR1 with AvrPphB results in cell death, whereas individual PBR1 fragments, or the auto-activating D496V NB-ARC mutation do not, signifying the necessity of all domains in *cis* (Jaiswal et al. 2023). *Pbr1* transcripts are significantly downregulated in the *mla6* mutant, but not in a *mla6 + bln1* double mutant, suggesting that the absence of *Bln1* in a *mla6* mutant background promotes normal *Pbr1* transcript accumulation (Figure 4). Another interpretation would simply be that wild-type *Mla6* is necessary for *Pbr1* expression. This pattern can also be classified under “dominant *Mla6*” in the *Mla6, Sgt1* model, indicating that *Sgt1* does not affect its expression. Nine of the 206 expressed genes with a consensus classification under the “*Mla6, Bln1* suppression” epistasis pattern were NLRs, with 7 uniquely downregulated in the *mla6* mutant, and 2 significantly upregulated (Figure 4, **Figure S5**). These observations indicate that NLR expression is interconnected and sensitive to gene-wise changes in the barley transcriptome, not only from NLRs, but from other immune-genes and pathogens.

Expression of NLRs in the masked category contrasts with suppression, in that a majority are up-regulated in the *bln1* single mutant, as compared to the *mla6 + bln1* double mutant (Figure 4); genes in this group are primarily clustered on chromosomes 1H, 3H, and 7H (Figure 2E). This suggests that in this case, the presence of *Bln1* is necessary to regulate gene expression, but control is released in its absence, except in combination a mutant *mla6.* Eight of the 206 expressed transcripts with a consensus classification under the “*Mla6, Bln1* masked” epistasis model were NLRs, with six uniquely up-regulated in the *bln1* mutant, and two significantly down-regulated (Figure 4). Considering that the *mla6* mutation abolishes ETI, while the *bln1* mutation enhances basal defense (Meng et al. 2009; Xu et al. 2015), we would expect that the masking pattern correlates with genes involved in the *Mla6*-based immune response. Indeed, genes under masked epistasis were associated with “translation” and “oxidation functions” (Figure 2D).

The second (*Mla6, Sgt1*) model uses the deletion of the NLR-type resistance gene *Mla6* (Halterman et al., 2001), but also takes advantage of an *in-frame* Lys-Leu deletion of *Sgt1*, which selectively disrupts race-specific resistance conditioned by the *Mla6*, *Mla7*, or *Mla12* alleles, but not *Mla1*, *Mla9*, *Mla10*, and *Mla13* (Chapman et al. 2021; Chapman et al. 2022). This model is designed to illustrate the interaction between *Mla6* and *Sgt1* when data from the double mutant is not available. It is not an epistasis model; however, it classifies gene expression using the wild-type progenitor as reference, and it quantifies the effects of each single-mutant. Previous studies have shown that SGT1 is required for MLA6-mediated generation of H_2_O_2_ and the hypersensitive reaction, and whole-plant resistance to *AVR_a6_* containing *Bh* isolates (Chapman et al. 2021; 2022). We separated these effects into dominant (when changes in expression are only observed in one of the single mutants; predominant (when there is effect of both genes but one of them is larger than the other; and equal effects (when both genes affect equally the gene expression). By applying these parameters, we could observe that most genes were under equal effects of *Mla6* and *Sgt1* or dominant effects from *Mla6*. This was similar to what was observed from the epistasis model between *Mla6* and *Bln1* (symmetric, followed by masking epistasis). As both *Mla6* and *Sgt1* mutations disrupt resistance to powdery mildew, a large overlap in the number of genes with equal effects from these genes is expected. Physical interaction between the proteins encoded by these genes has been also demonstrated, which is weakened with the mutated SGT1_ΔKL308-309_ protein, causing a reduction in MLA6 protein accumulation, and hindering resistance (Chapman et al. 2022). The GO terms “vesicle-mediated transport”, “S/T kinase amino acid” and “carbohydrate metabolism” were associated with the equal effects of *Mla6* and *Sgt1*, which suggests the presence of cellular functions that depend on a common transcriptional pathway, where both genes interact at the physical level.

Chromosomal location is postulated as a possible mechanism of action for the patterns observed with the gene effect models. Most of the gene effect patterns were enriched in chromosome regions as telomere-proximal (TP) and gene-rich interior (GRI). Trans-activation of gene expression has been associated with gene location (Cook and Marenduzzo 2018). Aspects such as genome accessibility and recruitment of transcriptional machinery through *cis*-regulatory elements may contribute to the association of gene effect patterns with genome location. For example, the relationship between histone modifications H3K27me3 and H3K27me1, has been associated with the determination of the TP and GRI regions, contributing to accessibility of the barley epigenome (Baker et al. 2015). We hypothesize that differential recruitment and activation of transcription factors is a key mechanism that regulates gene expression under the proposed gene effect models, although more in-depth study is necessary. In *Mla6*-based disease resistance, protein interactions have been demonstrated between the MLA6 protein and several transcription factors including WRKY, MYB, basic helix-loop-helix (bHLH) and homeobox (HB) (Shen et al. 2007; Chang et al. 2013; Velásquez-Zapata et al. 2021; Velásquez-Zapata et al. 2022). These interactions point to a direct mechanism of transcriptome regulation that can be modified by the loss-of-function mutations evaluated in the current report.

### Co-expression between Bh effectors and fungal development

On the pathogen side, we did not find evidence of epistasis, but that the co-expressed *Bh* infection transcriptome was concordant to disease phenotype. We constructed a *Bh* co-expression network and then associated it with infection kinetics traits via WGCNA (Langfelder and Horvath 2008; Figure 5 and **S1 Data**). This approach was effective to classify *Bh* gene expression, as the number of clusters and their separation were well defined. Early stages were associated with GO terms “metabolism”, “translation” and “metallopeptidase activity” (Aguilar et al. 2016; Thilini Chethana et al. 2021), whereas later stages were associated with “RNA binding”, “GTPase activity” and “proteasome” (Lambertucci et al. 2019; Pennington et al. 2019; Bauer et al. 2021).

Secreted proteins were distributed among all co-expression clusters, with significant enrichment in *Bh*3 (brown; 725 genes), *Bh7* (red; 246 genes) and *Bh*8 (turquoise; 1921 genes). When associations with traits was evaluated, we generally observed that the *Bh*3 (brown; 725 genes), *Bh*4 (green; 420 genes), *Bh*5 (grey; 2091 genes) and *Bh*9 (yellow; 613 genes) clusters were associated with attached conidiospores while *Bh*8 (turquoise; 1921 genes) linked with hyphal indexes 1-3, indicating that accumulation of transcripts encoding secreted proteins is aligned with the development of *Bh* structures and that these proteins play different roles over time. Finally, previous functional characterization of *Bh* effectors provided another layer of information for our co-expression analysis (Zhang et al. 2012; Pliego et al. 2013; Ahmed et al. 2015; Ahmed et al. 2016; Li et al. 2021; Yuan et al. 2021; Pennington et al. 2019). For example, effectors with associations to attached conidiospores and appressoria, including BLGH_00280 (CSEP0079; *Bh8*, turquoise), BLGH_04692 (CSEP0128; *Bh3*, brown), BLGH_01483 (CSEP0420; *Bh7*, red) and BLGH_00600 (CSEP0422; *Bh*8, turquoise) were also found to be involved in early fungal aggressiveness (Aguilar et al. 2016). In addition, BLGH_06477 (CSEP0041; *Bh9*, yellow) was previously found to be highly expressed in the appressorial germ tube (Pham et al. 2019).

Our analysis of the *Bh* transcriptome indicates an important role of the effectors associated with hyphal indexes as well, which were mostly classified in the *Bh8* (turquoise) module. This cluster comprises most of the reported AVR candidates and other effectors that have some functional characterization. Li et al. (2021) screened ∼100 *Bh* effector candidates and found that BLGH_07004 (CSEP0139) and BLGH_06939 (CSEP0182) suppressed BAX-induced programmed cell death in *Nicotiana benthamiana* and in barley. These two were found to be significantly associated with hyphal indexes 2 and 3, also known as elongating secondary hyphae (ESH), visual indicators of functional haustoria during development (Ellingboe 1972). *AVR_a6_* was also classified in the *Bh8* cluster. Interestingly, while BLGH_00709 (CSEP0254) has been reported as an *AVR_a6_* candidate*, Bh* DH14 also contains two near-identical copies, BLGH_00708 and BLGH_07091 (Bauer et al. 2021). In *Bh* 5874, only BLGH_0078 is expressed, and not BLGH_00709 or BLGH_07091. This suggests that different copies of *AVR_a6_* are expressed in an isolate-specific manner.

## DATA AVAILABILITY

Strains and plasmids are available upon request. Supplemental files available at FigShare. S1 Data contains count data for microscopic quantification of fungal structures. S2 Data contains the differentially expressed genes from the barley and *Bh* co-transcriptomes between wild-type and mutant genotypes. S3 Data contains the gene effect classifications per model and timepoint. S4 Data contains the enrichment of gene effect gene lists in chromosomal locations. S5 Data contains module and gene associations between the *Bh* co-expression network and phenotypic traits. Figure S1 contains GSE101304 sequencing experimental design. Figure S2 contains sequence and expression analysis of the *mla6* and *bln1* mutations. Figure S3 contains PCA plots of the barley-*Bh* transcriptome counts separated by species. Figure S4 contains GO enrichment analysis of the *Bh* co-expression network. Figure S5 contains graphs of all barley NLR expression patterns on CI 15161 and derived mutants from 0 to 48 HAI. S1 Text outlines the *Mla6, Bln1* and *Mla6, Sgt1* gene effect models. Infection-time-course RNA-Seq datasets are available at the NCBI-Gene Expression Omnibus (GEO) database accession number GSE101304 (https://www.ncbi.nlm.nih.gov/gds/?term=GSE101304). Supporting code for the analyses can be found in the GitHub repository https://github.com/Wiselab2/Epistasis.

## SUPPLEMENTAL FILES

Figure S1. GSE101304 sequencing experimental design.

Figure S2. Gene mutation characterization.

Figure S3. PCA of the barley-Bh transcriptome counts separated by species.

Figure S4. GO enrichment analysis of the Bh co-expression network.

Figure S5. Barley NLR expression patterns.

S1 Data. Count data for quantification of fungal structures.

S2 Data. DEG from the barley-*Bh* transcriptome between wild-type and mutant genotypes.

S3 Data. Barley gene effect classification per model and timepoint.

S4 Data. Enrichment of barley gene effect gene lists in chromosomal locations.

S5 Data. Association between the *Bh* co-expression network and infection kinetics.

S1 Text. Gene effect models.

## ACKNOWLEDGMENTS

The authors thank Gregory Fuerst for marker assisted selection and PCR-based sequence verification of the *mla6, bln1* and *mla6 + bln1* mutants, Stefan Kusch for the most up-to-date *Bh* DH14 genome assembly and annotation, Carsten Pedersen for sharing *Bh* CSEP annotations, and Matt Moscou for assistance with mapping of *Bln1* in the Morex v3 assembly. Research supported in part by National Science Foundation - Plant Genome Research Program grant 13-39348, USDA-National Institute of Food and Agriculture (NIFA) grant 2020-67013-31184 and USDA-Agricultural Research Service projects 3625-21000-067-00D and 5030-21220-068-000-D to RPW. The funders had no role in study design, data collection and analysis, decision to publish, or preparation of the manuscript. Mention of trade names or commercial products in this publication is solely for the purpose of providing specific information and does not imply recommendation or endorsement by the USDA, NIFA, ARS, or the National Science Foundation. USDA is an equal opportunity provider and employer.

## AUTHOR CONTRIBUTIONS

Designed the research: VVZ, RPW.

Performed research: VVZ, PS, AC.

Contributed new analytic/computational tools: VVZ, SS.

Analyzed data: VVZ, PS, SS, RPW.

Wrote the paper: VVZ.

Edited the paper: VVZ, RPW.

